# CEP162 deficiency causes human retinal degeneration and reveals a dual role in ciliogenesis and neurogenesis

**DOI:** 10.1101/2021.11.23.469779

**Authors:** Nafisa Nuzhat, Kristof Van Schil, Sandra Liakopoulos, Miriam Bauwens, Alfredo Duenas Rey, Stephan Käseberg, Melanie Jäger, Jason R. Willer, Jennifer Winter, Hanh Truong, Nuria Gruartmoner, Mattias Van Heetvelde, Joachim Wolf, Robert Merget, Sabine Grasshoff-Derr, Jo Van Dorpe, Anne Hoorens, Heidi Stöhr, Luke Mansard, Anne-Françoise Roux, Thomas Langmann, Katharina Dannhausen, David Rosenkranz, Karl Martin Wissing, Michel Van Lint, Heidi Rossmann, Friederike Häuser, Peter Nürnberg, Holger Thiele, Ulrich Zechner, Jillian N. Pearring, Elfride De Baere, Hanno J. Bolz

## Abstract

Defects in primary or motile cilia result in a variety of human pathologies, and retinal degeneration is frequently associated with these so-called ciliopathies. We show that homozygosity for a truncating variant in CEP162, a centrosome and microtubule-associated protein required for transition zone (TZ) assembly during ciliogenesis and neuronal differentiation in the retina, causes late-onset retinitis pigmentosa in 2 unrelated families. The mutant CEP162-E646R*5 protein is expressed and properly localized to the mitotic spindle but missing from the basal body in primary and photoreceptor cilia. This impairs recruitment of TZ components to the basal body and corresponds to complete loss of CEP162 function at the ciliary compartment, reflected by delayed formation of dysmorphic cilia. In contrast, rescue of increased cell death in the developing mouse retina after shRNA knockdown of *Cep162* by expression of CEP162-E646R*5 indicates that the mutant retains its role for retinal neurogenesis. Human retinal degeneration thus results from specific loss of ciliary CEP162 function.

**Graphical abstract:** 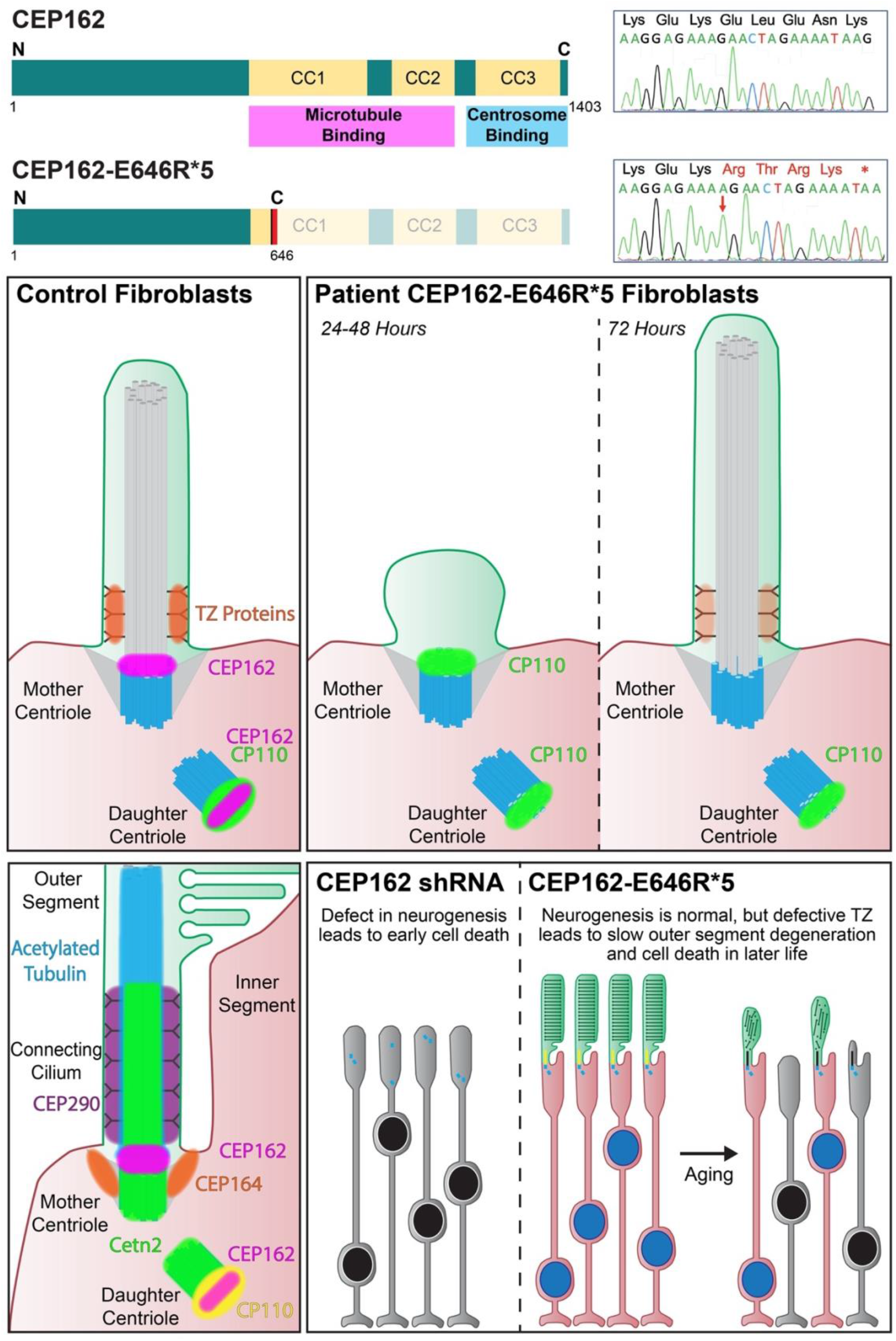

## Introduction

Primary cilia are sensory organelles that protrude from the cell surface to detect extracellular signals that regulate cellular physiology. Cilia are highly dynamic microtubule-based organelles that are nucleated from the mother centriole and assembled and disassembled in each round of the cell cycle. The complex process of cilium assembly, maintenance, and disassembly is estimated to require thousands of genes, with nearly 700 transcripts confirmed to be localized to the cilium (1) and many more implicated in ciliary function. Dysfunction of ciliary genes and proteins is associated with a wide range of human disorders. These ciliopathies can involve virtually any organ, with particularly frequent affection of the retina. Proteins with established roles in cilia may also participate in other cellular processes. For example, OFD1, a centrosomal protein of the basal body implicated in isolated retinitis pigmentosa (RP) and different syndromes (2, 3), is also involved in chromatin remodeling (4) and cell cycle progression (5).

*CEP162* was originally cloned from quail retinal neurons and designated quail neuroretina 1, *QN1* (6). Inhibition of QN1 during retinal development lead to defective mitosis and differentiation suggesting that it is involved in withdrawal from cell cycle, so QN1 was postulated to play a role in neuronal quiescence (7, 8). QN1 is orthologous to human KIAA1009/CEP162 (9), a protein that binds to microtubule spindles during mitosis and localizes to the distal ends of centrioles in post-mitotic cells (8, 10). Loss of CEP162 has been shown to arrest ciliogenesis at the stage of TZ assembly and to cause a ciliopathy phenotype in zebrafish (10). Despite its importance for cilia formation, no pathogenic *CEP162* variants have been reported in human disease, and its role in the retina has remained unknown.

Here, we show that homozygosity for a frameshift variant in *CEP162* causes late-onset RP in 2 unrelated families. We find that the truncated CEP162 is unable to localize to the basal body. Absence of CEP162 from the primary cilium results in a loss of some TZ components and delayed formation of dysmorphic cilia. However, truncated CEP162 maintains its capability to bind microtubules and localizes to the mitotic spindle, suggesting that it could retain function in neuronal cell division. Indeed, we find that increased cell death in the developing mouse retina with loss of *Cep162* expression can be restored not only by full-length, but also truncated CEP162 which is unable to localize to basal body in these cells. Thus, specific loss of CEP162 function at the primary cilium is likely the primary cause of the late-onset human retinal ciliopathy.

## Results

### Late-onset retinitis pigmentosa (RP) in 2 unrelated patients

*Patient 1*. Loss of visual acuity was noted at 56 years of age. At 60 years, RP was diagnosed. Night blindness had presumably pre-existed for years. There is now noticeable photophobia. At the age of 67 years, refraction revealed hyperopia and astigmatism in both eyes with +6.25 Sphere (sph), −1.25 Cylinder (cyl)/71° in the right eye and +2.75 sph, −1.00 cyl/58° in the left eye. Color fundus photography, blue light autofluorescence and spectral domain optical coherence tomography (OCT) were compatible with RP (pale optic disc, narrow vessels, bone spicule pigmentation, thinning of the outer nuclear layer; Figure 1A-F). Best corrected visual acuity (BCVA) was 20/500 for both eyes (Snellen chart), and visual fields were constricted. There was *synchysis scintillans* in the right vitreous. *Patient 2*. Loss of visual acuity was reported since the 5^th^ decade. At the age of 69 years, RP was diagnosed. BCVA was light perception (OD) and 4/10 (OS) and visual fields were constricted <10°. Color fundus photography showed a pale optic disk, narrow vessels with sheathing and limited bone spicule pigmentation. Fluorescein angiography displayed strong atrophy of the outer retina and few intraretinal pigment migrations. OCT showed absence of the outer retina, except in the fovea. On blue light autofluorescence, a Bull’s eye aspect was observed. A pattern electroretinogram (ERG) showed an absent response (OD) and reduced macular function (OS) (Figure 1G-N). A full-field ERG could not be conducted. Patient 2 was followed at the diabetes clinic for 30 years; no islet cell autoantibodies were detected. There was a low normal production of C-peptide (0.20 nmol/L. Ref: 0.29-0,99 nmol/L). He received medication for chronic renal insufficiency due to diabetic nephropathy. Abdominal ultrasound and CT showed no kidney abnormalities (Supplemental Figure 1), but a lipomatous aspect of the pancreas. Patient 2 suffered from coronary main stem stenosis (50-59%, non-invasive treatment) and died at the age of 74.

**FIGURE 1.**
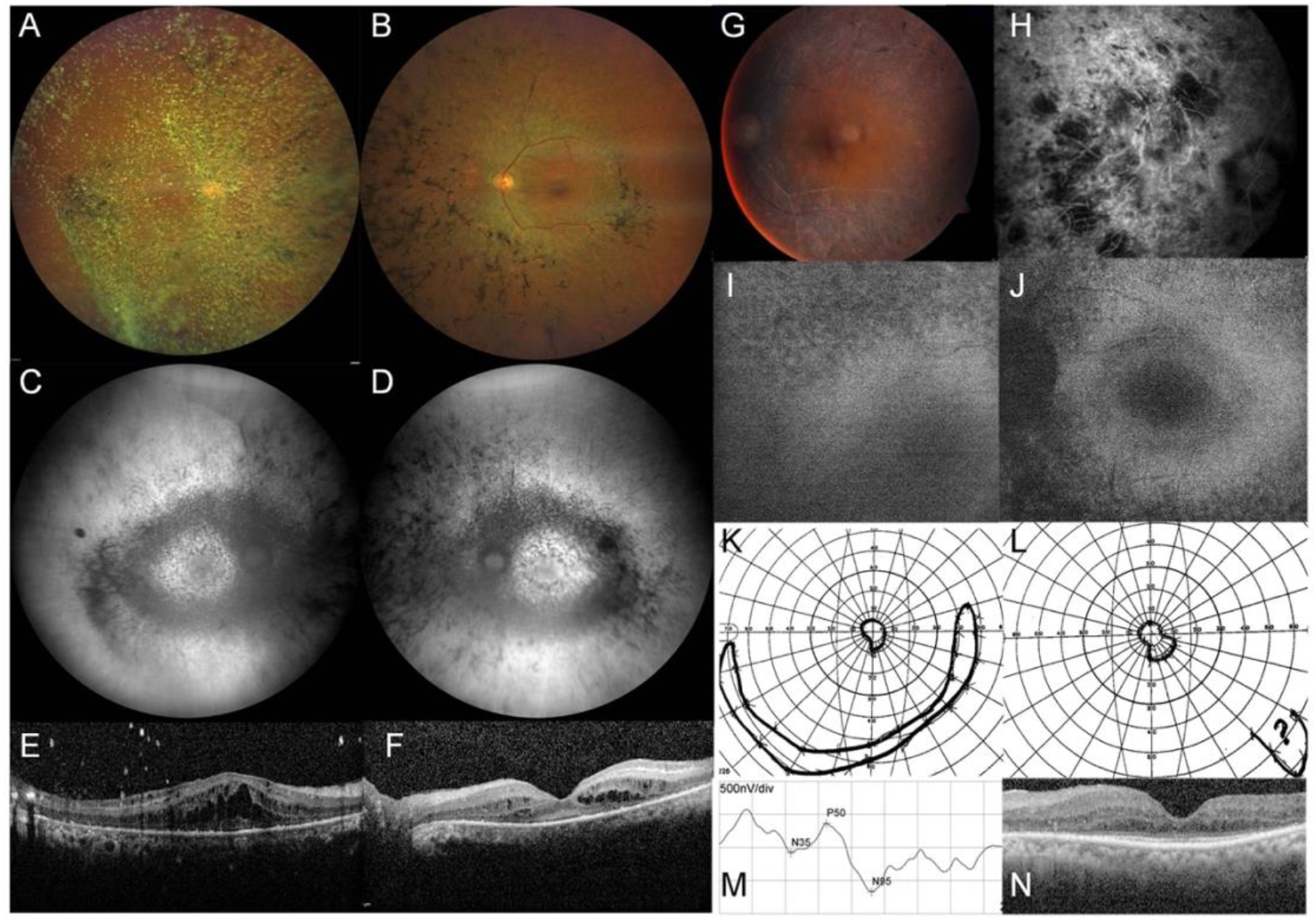
Ophthalmological data of Patient 1 (A-F) and Patient 2 (G-N) with RP. **A** Right eye (OD, left column) and **B** left eye (OS, right column) with pale optic disc, narrow vessels and bone spicule pigmentation on color fundus photography. **C, D** Granular decreased autofluorescence throughout the posterior pole on blue light autofluorescence. **E, F** SDOCT: Cystoid spaces in the inner and outer nuclear layer and thinning of the outer nuclear layer with sparing of the central fovea. **G** Color fundus photography of the left eye: Pale optic disc, narrow vessels with pronounced sheathing giving a white (pseudo-thrombotic) aspect, bone spicule pigmentation. **H** Fluorescein angiography of the right eye: Strong atrophy of the outer retina and few intraretinal pigment migrations. Autofluorescence from **I** right eye and **J** left eye: Bull’s eye aspect of the macula. Goldmann visual fields for **K** left eye and **L** right eye: constriction of <10°. **M** Pattern ERG of left eye: Reduced macular activity (visual acuity 3/10). No responses for the right eye. **N** OCT: Absence of the outer nuclear layer beyond the macula.

### Homozygous *CEP162* frameshift variant in both unrelated RP patients

Patient 1 and Patient 2 were both born to consanguineous parents in 2 unrelated Moroccan families (Figure 2A). *Patient 1*. The grandparents of Patient 1, a 68-years old male, are 2^nd^ degree cousins. No pathogenic variant was identified in Patient 1 by targeted NGS of 204 known IRD genes. WES generated 16.3 Gb of sequence, covering 97.9% of target sequence >30x. Among 701 rare variants, 41 were homozygous, further narrowed to 25 by applying an MAF of <1%, excluding artifacts and filtering with ROH. 24 variants were disqualified as RP-causing due to 1 or several of these reasons: a) Gene associated with an unrelated disease, b) homo- or hemizygous individuals in gnomAD, c) variant predicted as neutral/benign. 1 variant remained, a 1-base pair insertion in exon 15 of *CEP162*, NM_014895.3:c.1935dupA, causing a frameshift and premature nonsense codon (p.(Glu646Argfs*5), subsequently designated as E646R*5; Figure 2B, D). In the other annotated *CEP162* isoform, the insertion corresponds to NM_001286206.1:c.1707dup, p.(Glu570Argfs*5).

**FIGURE 2.**
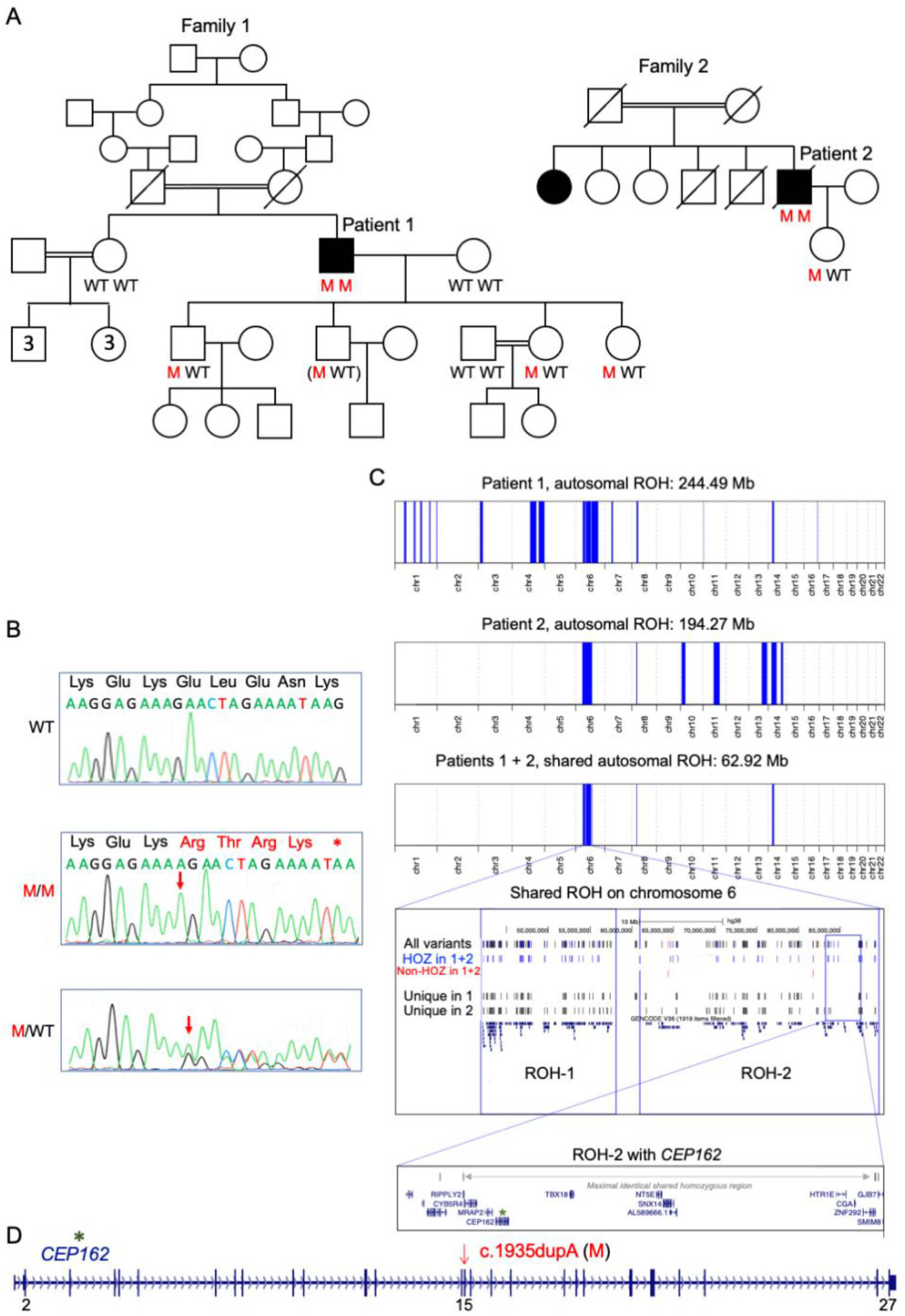
Homozygous *CEP162* frameshift mutation causes RP in 2 unrelated Moroccan families. **A** Pedigrees with individuals who were available for genotyping of the c.1935dupA (p.(E646R*5)) in *CEP162*. M, mutation; WT, wild type. **B** Electropherograms of an individual with WT sequence (upper panel), Patient 1 (middle; homozygous 1-bp insertion with frameshift and premature termination codon), and a heterozygous carrier (bottom; all children of the patients). **C** ROH on chromosome 6, comprising *CEP162*, shared by Patient 1 and Patient 2. **D** Scheme of *CEP162* gene (to scale). Vertical bars: exons (pathogenic variant in exon 15).

*Patient 2.* His older affected sister, living in North Morocco, was not available for testing. WES in the proband generated 40.6 million reads, with 99.0% of reads mapping to the target sequences, providing an average coverage of >30x. Assessment of 275 RetNet genes did not reveal any (likely) pathogenic variants. An exome-wide analysis revealed 1,298 rare variants (MAF <1%), 190 of which were homozygous. These were reduced to 58, filtering with ROH (Figure 2C, Supplemental Table 1), and ultimately to the *CEP162* c.1935dupA variant, based on the aforementioned criteria. *Patient 1 & 2*. Segregation analysis in both families supported a pathogenic nature of the truncating *CEP162* variant (Figure 2A-B; Supplemental Figure 2) which was absent from gnomAD (v.2.1.1). No homozygous *CEP162* LoF variants were found in gnomAD. The variant was neither found in 70 RP patients from North Africa (incl. 43 from Morocco) by targeted testing, nor in WES data from 1,184 IRD cases. The largest ROH in both Patient 1 and 2 is located on chromosome 6 (Figure 2C; Supplemental Table 1) and includes the *CEP162*:c.1935dupA variant, putting it forward as a potential founder allele (Figure 2C).

### Mutated c.1935dupA *CEP162* mRNA escapes nonsense-mediated decay in patient fibroblasts, allowing for expression of truncated CEP162 protein

Fibroblasts from Patient 1 were compared to control human dermal fibroblasts (Hdfa). qRT-PCR analysis revealed that *CEP162* transcript levels were significantly reduced in patient fibroblasts (Supplemental Figure 3A). *CEP162* mRNA levels significantly increased upon serum starvation, suggesting that the transcriptional regulation of the *CEP162* gene remained unaffected (Supplemental Figure 3A). Anisomycin treatment was used to determine whether the patient *CEP162* transcript undergoes nonsense-mediated decay (NMD). Increased expression of *CEP162* after anisomycin treatment was observed in both patient and control cells (Supplemental Figure 3B), suggesting that a basal level of *CEP162* transcript physiologically undergoes NMD.

To determine whether the truncated CEP162-E646R*5 protein is expressed in patient fibroblasts, we immunoblotted control and patient fibroblast lysates with an antibody that recognizes the N-terminus of CEP162 prior to the truncation. Patient lysates only had a band at the predicted size of truncated CEP162 (~75 kDa) compared the full-length CEP162 (~160 kDa) band in control lysates (Supplemental Figure 3C). In addition, probing with the C-terminus anti-CEP162 antibody produced a full-length CEP162 band in control lysate, but no corresponding or truncated band in the patient lysate (Supplemental Figure 3C). We conclude that despite the lower levels of *CEP162* mRNA, the residual transcript does not undergo complete NMD, resulting in expression of truncated CEP162-E646R*5 protein in patient cells (Supplemental Figure 3D).

### CEP162-E646R*5 mutant protein binds microtubules but is unable to associate with centrioles or CEP290

Human CEP162 comprises 1,403 amino acids with 3 coiled-coil (CC) stretches in its C-terminus: CC1 (residues 617-906), CC2 (957-1,121) and CC3 (1,167-1,386). CEP162 associates with microtubules through its CC1/CC2 domains and with centrioles through its CC3 domain (10). The c.1935dupA mutation results in early truncation of the protein after the first 29 amino acids of the CC1 domain (Figure 3A). Truncated FLAG-CEP162-E646R*5 produced a ~80 kDa band compared to the full-length FLAG-CEP162 band at ~165 kDa (Figure 3B, Supplemental Figure 4A). We performed a microtubule binding assay to determine whether the CEP162-E646R*5 mutant protein binds microtubules. Immunoprecipitated FLAG-tagged human CEP162 full-length or E646R*5 mutant protein were co-pelleted with taxol-stabilized microtubules. Figure 3C shows both full length FLAG-CEP162 and truncated FLAG-CEP162-E646R*5 were found in the microtubule pellets suggesting that the residual CC1 domain is sufficient to bind microtubules.

**FIGURE 3.**
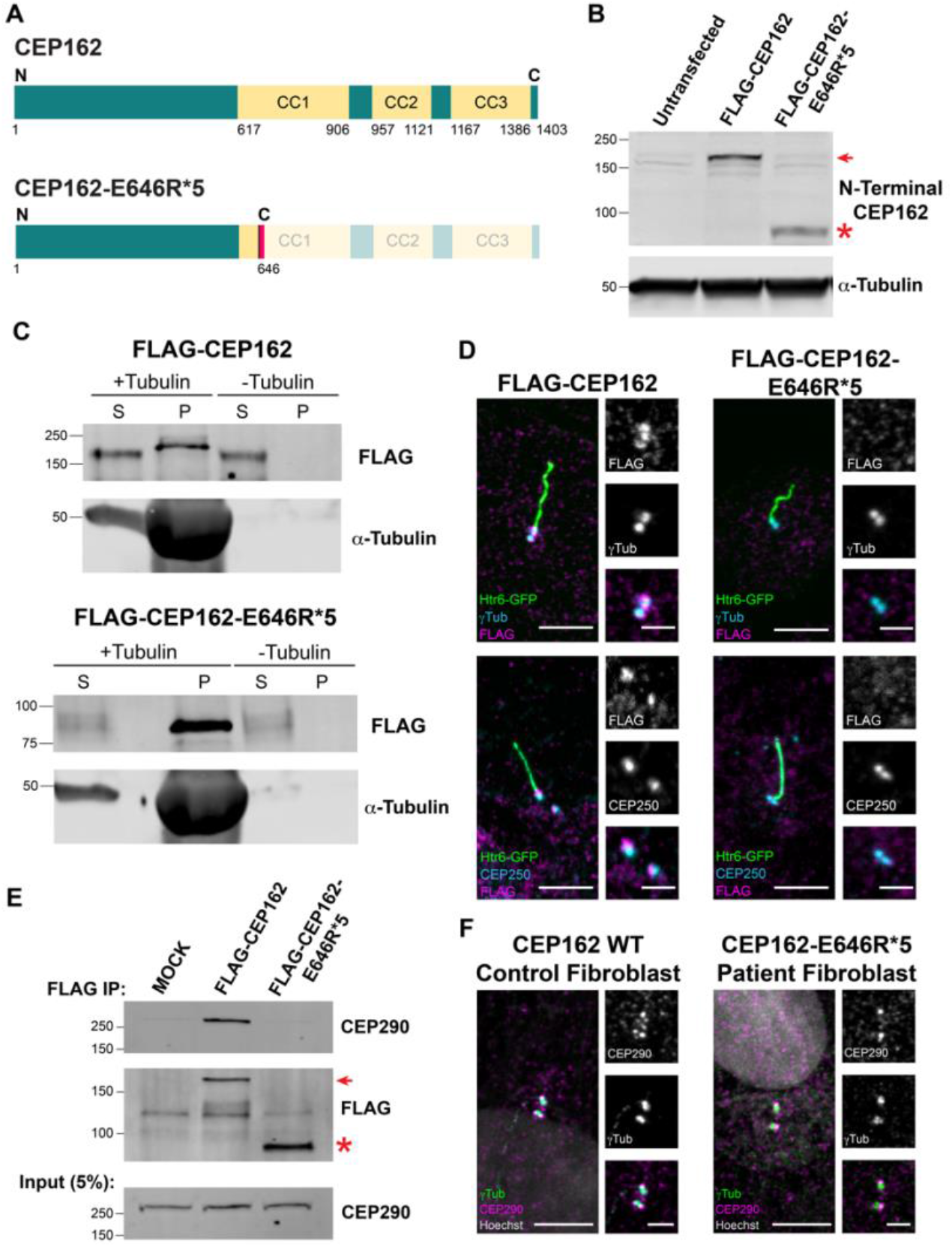
Effect of E646R*5 mutation on CEP162 protein expression and localization. **A** Scheme of human CEP162 protein with 3 C-terminal coiled-coil (CC) domains and the truncated CEP162-E646R*5 mutant protein. The amino acid residues are given for each scheme. **B** Western blot of 239T cell lysates from untransfected control, transfected FLAG-CEP162 (~165 kDa, red arrow) and FLAG-CEP162-E646R*5 (~80 kDa, red asterisk). Blots were probed for CEP162 to detect expressed protein and α-tubulin as a loading control. **C** Microtubule binding assay. Purified FLAG-CEP162 or FLAG-CEP16-E646R*5 was co-pelleted with taxol-stabilized microtubules. CEP162 was probed by anti-FLAG antibodies and pelleted microtubules were detected by anti-αTubulin antibodies. S, supernatant; P, pellet. **D** Serum-starved IMCD3 cells co-expressing Htr6-GFP and FLAG-CEP162 or FLAG-CEP162-E646R*5. Transfected cells were identified by GFP (green) fluorescence in the cilium. FLAG (magenta) was co-immunostained with either γ-tubulin (cyan, basal bodies) or CEP250 (cyan, proximal-end centriolar protein). **E** FLAG immunoprecipitation from Mock, FLAG-CEP162 or FLAG-CEP162-E646R*5 transfected 293T cells lysates. Endogenous CEP290 pulls down with full length FLAG-CEP162, but not MOCK or FLAG-CEP162-E646R*5. **F** CEP290 (magenta) immunostaining in control and patient fibroblasts co-stained with γ-tubulin (green). Higher magnification images of the cilium/basal body and corresponding staining are shown to the right. Scale bars, 5 μm and 2 μm.

To determine whether this truncated mutant protein associates with the centrioles, we co-transfected the FLAG-tagged CEP162 constructs with the ciliary marker Htr6-GFP into IMCD3 cells and serum-starved for 24 hours to ciliate the cells (Figure 3D). Full-length FLAG-CEP162 was present at the basal bodies of Htr6-GFP-positive cilia where it co-localizes with γ-tubulin. In addition, co-staining with a proximal-end centriole protein, CEP250, showed proper localization of FLAG-CEP162 to the distal end of the centrioles. In contrast, the mutant FLAG-CEP162-E646R*5 was not detected at the basal body (Figure 3D).

It was previously shown that CEP162 interacts with CEP290 through its CC1/CC2 domain (10). To determine whether the mutant CEP162-E646R*5 protein interacts with endogenous CEP290, we performed FLAG immunoprecipitations from our transfected 293T cells expressing either full length or mutant CEP162. FLAG-CEP162-E646R*5 was unable to pull down CEP290; however, CEP290 localization at the centrioles was normal in the patient fibroblasts (Figure 3E, F). Also, no difference in CEP290 protein levels was observed in patient fibroblasts compared to controls (Supplemental Figure 5).

### Patient fibroblasts have delayed ciliation

CEP162 is present at the centrioles throughout the cell cycle, and its microtubule-binding activity is believed to direct its localization to the mitotic spindle during cell division (10). Patient fibroblasts expressing the CEP162-E646R*5 truncated protein had a normal growth rate and no aneuploidy was detected in 30 metaphases. Consistent with our result showing that microtubule binding is retained by truncated CEP162-E646R*5, CEP162 staining decorated the microtubule spindles of dividing cells in the patient fibroblasts similar to controls (Figure 4A). This result suggests that truncated CEP162-E646R*5 localization and function is normal during fibroblast mitosis, which is expected since CEP162’s role in post-mitotic quiescence has only been described for neurons (7, 8).

**FIGURE 4.**
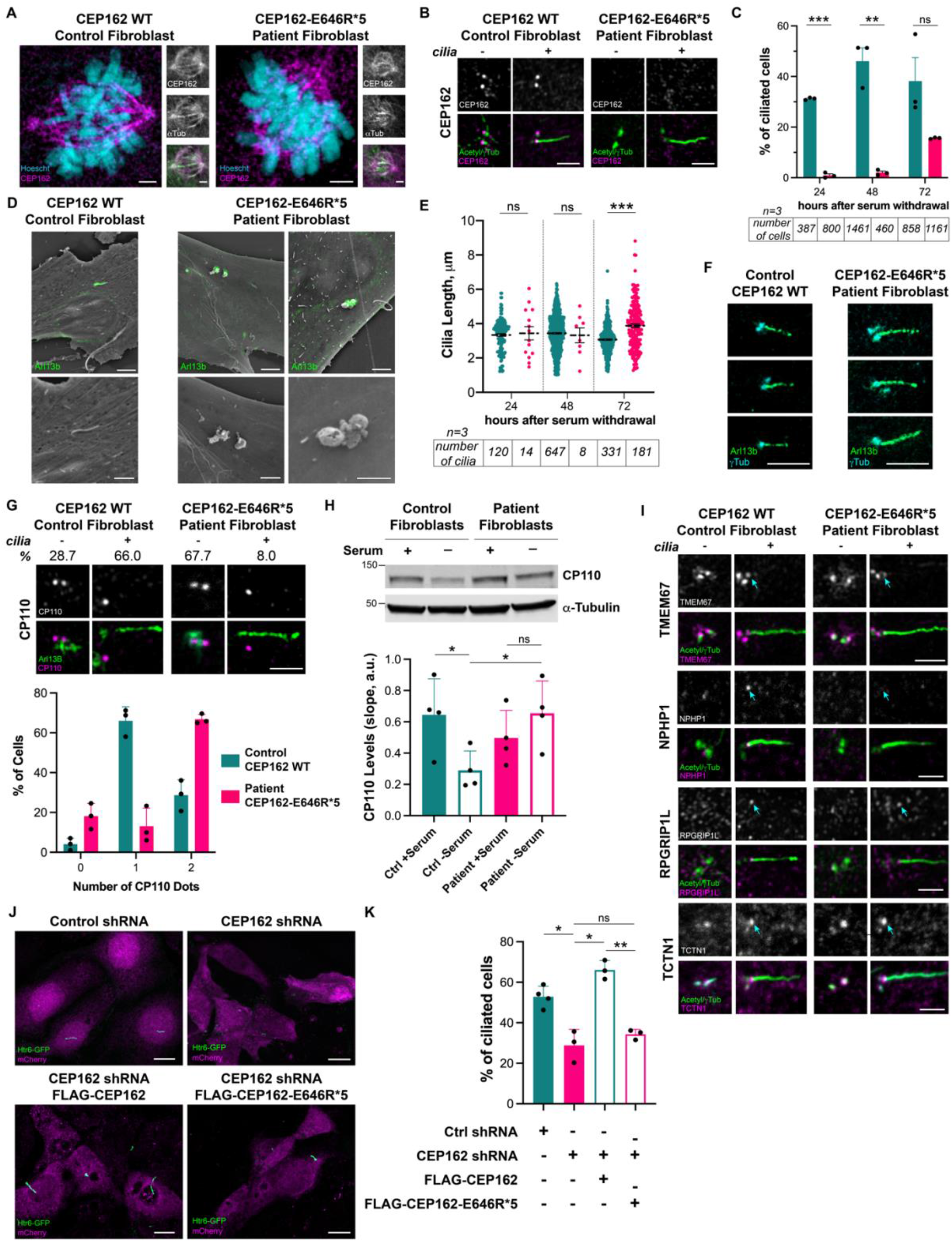
CEP162-E646R*5 localizes to the mitotic spindle in patient fibroblasts, but its absence from the basal body delays ciliation. **A** Control and patient fibroblasts co-stained for endogenous CEP162 (magenta) and α-tubulin show CEP162-E646R*5 protein at the mitotic spindle. Individual staining is shown to the right. Scale bars 2 μm. **B** CEP162 (magenta) immunostaining in control and patient fibroblasts co-stained with acetylated/γ-tubulin (Acetyl/γTub, green). Scale bar, 2 μm. **C** Percent ciliation in control and patient fibroblasts after 24, 48 or 72 hours after serum withdrawal was determined by Arl13b and γ-tubulin staining. ***p<0.000001, **p<0.001 **D** Correlative light and scanning electron microscopy were performed in control and patient fibroblasts. Cilia were identified by staining for Arl13b (green). Merged images are shown for the low scanning electron micrograph with magnified micrographs shown below. Scale bars, 4 μm and 2 μm. **E** Cilia length was measured for each cilium. ***p<0.0001 **F** Representative cilia from control and patient fibroblasts stained with Arl13b (green) and γ-tubulin (cyan). Scale bar, 5 μm. **G** Immunostaining of CP110 (magenta) in control and patient fibroblasts, with (+) and without (-) cilia. Percent observed below. Bar graph quantifying CP110 dots in control and patient fibroblasts. **H** Representative Western blot from control and patient fibroblast lysates after 24 hours with (+) or without (–) serum. Blots were probed for CP110 and α-tubulin for a loading control. Bar graph quantifying relative levels of CP110 in control and patient fibroblasts +/– serum from 4 independent experiments. *p<0.035 **I** Immunostaining of transition zone proteins TMEM67, RPGRIP1L, NPHP1 and TCTN1 (magenta) in control and patient fibroblasts, +/– cilia. In all panels, cilia/basal bodies were co-stained with acetylated/γ-tubulin (Acetyl/γTub, green), and cyan arrows mark transition zone. Scale bars, 2 μm. **J** Representative immunofluorescence images showing cilia (Htr6-GFP, green) in control or CEP162 shRNA targeted cells (mCherry, magenta) as well as CEP162 shRNA targeted cells co-expressing either full-length FLAG-CEP162 or FLAG-CEP162-E646R*5. Scale bars, 10 μm. **K** Percent ciliation in control or CEP162 shRNA expressing IMCD3 cells. Ciliation was rescued by expression of the full-length FLAG-CEP162, but not FLAG-CEP162-E646R*5. *p<0.019, **p<0.0019

As for ciliary localization, we found that CEP162 co-localizes with γ-tubulin-positive centrioles in control fibroblasts but was not detected in patient fibroblasts (Figure 4B). While control fibroblasts produced cilia within 24 hours, very few cilia in patient fibroblasts were observed before 72 hours serum withdrawal (Figure 4C). We used correlative light and scanning-electron microscopy (CLSEM) to examine the structure of the arrested cilia in the patient fibroblasts and compared them to normal cilia produced in controls. While control fibroblasts had Arl13b-positive primary cilia of normal length, patient fibroblasts only produced Arl13b-positive blebs on their surface (Figure 4D). Additionally, polyglutamylated tubulin staining was not observed in the stalled patient cilia indicating ciliogenesis is halted prior to axoneme extension in patient fibroblasts (Supplemental Figure 4B). Together, this suggests that the ciliary membrane has fused with the plasma membrane, resulting in a bulbous bleb on the cell’s surface.

By 72 hours after serum withdrawal, the number of cilia produced in the patient fibroblasts was not significantly different from controls (Figure 4C). Ciliary length was measured for every cilium imaged at the 3 timepoints. At 72 hours, the patient cilia were significantly longer than the controls (p>0.0001, Figure 4E, F). Together, our data suggest that in patient fibroblasts, ciliogenesis is paused before axoneme extension, but these cells can ultimately overcome the loss of CEP162 at the centrioles and produce cilia. Accumulation of ciliary membrane before axoneme elongation could affect the final length of the cilium in the patient fibroblasts.

### Persistence of CP110 at the mother centriole delays primary ciliogenesis in patient fibroblasts

To determine the stage at which ciliogenesis is paused in patient fibroblasts, we analyzed the localization of molecular components involved in cilia formation. First, maturation of the basal body appears to be normal as proteins involved in ciliary vesicle fusion (EHD1), distal appendages formation (CEP164) and IFT machinery recruitment (IFT88) were normal (Supplemental Figure 6A). Following ciliary vesicle formation, CP110, a distal-end centriole protein that prevents microtubule nucleation, is removed from the mother centriole, and degraded to enable proper axoneme elongation (11). This regulatory step was previously reported to be unaffected by loss of CEP162 in RPE1 cells (10). In patient fibroblasts 48 hours post serum withdrawal, quantification of CP110 dots revealed persistence of CP110 at the mother centriole of stalled cilia (Figure 4G). In addition, Western blot analysis for CP110 indicates that while levels normally decrease 50% upon serum starvation in controls, levels of CP110 remained unchanged in the patient fibroblasts 24 hours after serum withdrawal (Figure 4H; Supplemental Figure 5). CP110 removal is controlled by CEP164-mediated recruitment of the serine/threonine protein kinase, TTBK2, to the distal appendages (12, 13). TTBK2 phosphorylates multiple targets, such as CEP83 and MPP9 (11, 14), that are required for CP110 removal and ciliogenesis. We found both TTBK2 and its MPP9 substrate were normally localized in the patient fibroblasts (Supplemental Figure 6B). Although the molecules needed to remove CP110 from the mother centriole are present, this process is delayed by the loss of CEP162 at the basal body.

It was reported by Wang *et al*. (10) that exogenous expression of a C-terminally truncated CEP162, that maintains microtubule association but is unable to localize to centrioles, resulted in cilia that were abnormally long. In these experiments, C-terminally truncated CEP162 was found at the axoneme tip of elongated cilia where it ectopically recruited TZ components (10). Our data show that E646R*5-truncated CEP162 maintains microtubule binding during mitosis but is not localized to centrioles, suggesting it may behave similarly. To test this, we stained control and patient fibroblasts for 4 TZ proteins: TMEM67, NPHP1, RPGRIP1L, and TCTN1 (Figure 4I). We found normal localization of TCTN1, but TMEM67, NPHP1 and RPGRIP1L were not properly assembled at the ciliary base of patient fibroblasts, consistent with the complete loss of CEP162 (10). Importantly, we did not find mislocalization of any of these TZ components to the tip of the long patient cilia (Figure 4I), indicating that the CEP162-E646R*5 protein has lost its ability to recruit TZ proteins to cilia.

To confirm that CEP162-E646R*5 has no ciliary function, we performed shRNA knockdown of *CEP162* in IMCD3 cells, followed by rescue with either FLAG-tagged full-length CEP162 or CEP162-E646R*5. A single plasmid co-expressing shRNA and soluble mCherry was used to identify transfected cells and were co-transfected with an Htr6-GFP plasmid to label the cilia. Figure 4J shows images of the cilia in mCherry-positive cells when expressing the control or *CEP162* shRNA. *CEP162* knockdown resulted in reduced ciliation compared to control shRNA, similar to previous results (10). Expression of the FLAG-tagged full-length CEP162 rescued the loss of cilia due to *CEP162* knockdown; however, FLAG-CEP162-E646R*5 was unable to rescue (Figure 4J, K). These results support that expression of CEP162-E646R*5 protein does not retain any ciliary function.

### CEP162 is expressed in human retina and localizes to the basal body in mouse photoreceptors

We examined *CEP162* expression in single-cell transcriptional data of human neural retina (15). This revealed high expression in all retinal cell types, especially in ganglion cells (Supplemental Figure 7A). Immunostaining for CEP162 on human retinal sections shows expression throughout the human retina (Supplemental Figure 7B-F), similar to expression previously determined in chick retina (7). CEP162 is a centriolar protein; to determine its precise localization in photoreceptors, we used Cetn2-GFP transgenic mice that have fluorescently labelled centrioles (16). In photoreceptors, mother and daughter centrioles form 2 adjacent GFP dots, and the connecting cilium emerges as a GFP streak from the mother centriole. CEP162 staining is localized to the distal end of both mother and daughter centrioles (Figure 5A). Airyscan images were acquired to determine the precise localization of CEP162 in relation to several ciliary markers: Acetylated tubulin for the axoneme, CEP290 for the connecting cilium, CEP164 for the distal appendages, and CP110 for the daughter centriole (Figure 5B). CEP162 decorates the distal ends of each centriole at the base of the photoreceptor outer segment in wild-type mouse retinas.

**FIGURE 5.**
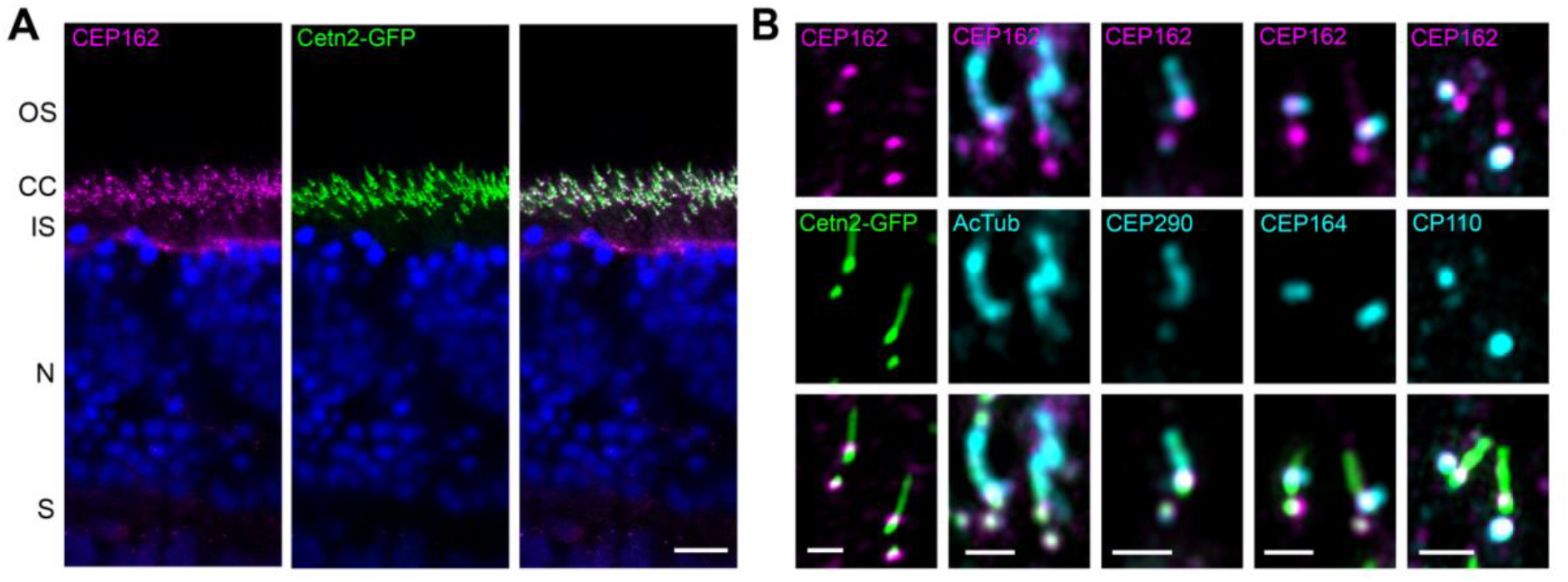
Endogenous CEP162 is localized to the distal-end of centrioles at the base of the photoreceptor outer segment in adult wild-type mouse retina. **A** Cetn2-GFP transgenic mouse retinal sections stained with an anti-CEP162 (magenta) antibody: CEP162 is localized at the photoreceptor connecting cilium, which is marked by GFP fluorescence (green). Scale bar, 10 μm. **B** High-resolution Airyscan images: CEP162 (magenta) staining decorates the distal ends of each centriole of the basal body at the base of the connecting cilium (Cetn2-GFP, green). Additional Airyscan images of CEP162 counterstained with multiple ciliary markers: Acetylated Tubulin (AcTub), CEP290, CEP164, and CP110 (cyan). Scale bar, 1 μm.

### CEP162-E646R*5 is unable to localize to the basal body in mouse photoreceptors but can rescue neuronal cell death

To determine the localization of truncated CEP162-E646R*5 mutant protein in photoreceptors, we employed *in vivo* electroporation (17) to express FLAG-tagged full-length and E646R*5-mutant *CEP162* constructs in mouse rods. We co-expressed Rho-mCherry to label the outer segment in transfected rods and stained retinas with anti-Centrin1 antibodies to label the connecting cilium in all photoreceptors. Figure 6A shows FLAG-CEP162 staining at the basal body directly below the connecting cilium in rods with Rho-mCherry-labelled outer segments. In contrast, FLAG-CEP162-E646R*5-mutant staining does not localize to the basal body of Rho-mCherry-labelled outer segments (Figure 6B). This suggests that the truncated CEP162-E646R*5-mutant protein is unable to properly localize to the centrioles in photoreceptors.

**FIGURE 6.**
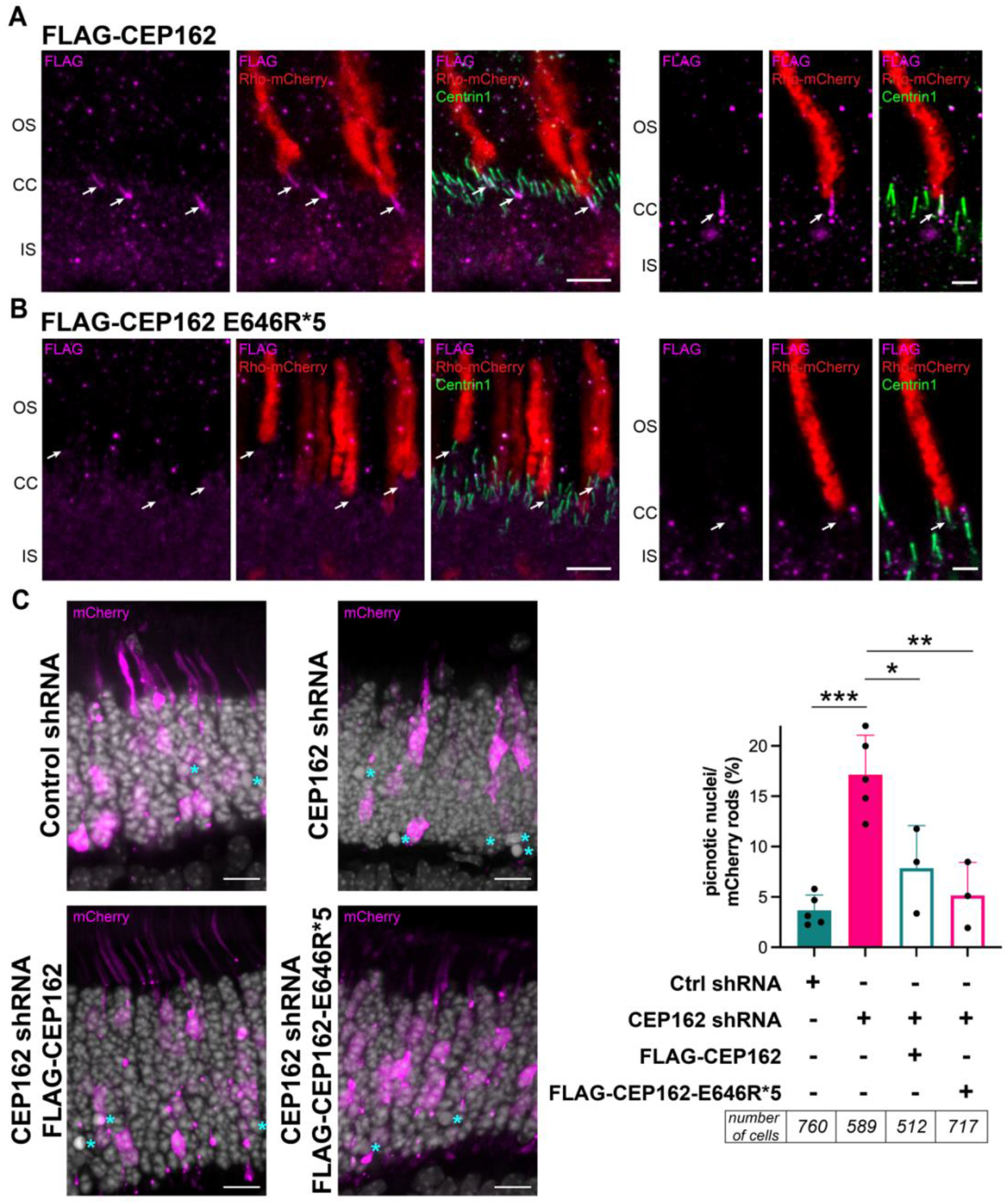
CEP162-E646R*5 mutant protein does not localize to centrioles in adult mouse rod photoreceptors but does participate in retinal neurogenesis. **A-B** Wild-type mouse rods were transfected with FLAG-CEP162 (**A**) or FLAG-CEP162-E646R*5 (**B**, magenta) and rhodopsin-mCherry (red) as a transfection marker. Centrin1 (green) immunostaining was conducted to label the connecting cilium. Arrows indicate the basal body of the transfected rods. Scale bars, 5 μm and 2 μm. **C** Developing mouse retina (P0-P1) were electroporated with control or CEP162 shRNA plasmids co-expressing a soluble mCherry (magenta), and CEP162 knockdown was rescued with either FLAG-CEP162 or FLAG-CEP162-E646R*5. Scale bar, 5 μm. Cell death was assessed by counting pycnotic nuclei labeled with DAPI (grey, cyan asterisk) and quantified for a minimum of 3 separate animals. ***p<0.0007, **p<0.0054, *p<0.036.

Using a retroviral antisense knockdown strategy in developing chick eyes, CEP162 (aka QN1) was reported to play a key role in differentiation of retinal neurons (7). We found that CEP162-E646R*5 maintains microtubule binding at the mitotic spindle in fibroblasts, so it could possibly function in neuronal development. To test this, we *in vivo*-electroporated *CEP162* shRNAs during retinal development and rescued with full-length or truncated *CEP162* constructs. Figure 6C shows representative retinal sections from P15 mice expressing a single plasmid co-expressing control or *CEP162* shRNA and soluble mCherry to identify electroporated rods. Pycnotic nuclei were counted within each electroporated area and normalized to the number of mCherry positive rods. Knockdown of *CEP162* increased cell death ~4-fold compared to controls, which was rescued by expression of either full-length or CEP162-E646R*5 (Figure 6C).

In conclusion, our data suggest that CEP162-E646R*5 retains microtubule binding at the mitotic spindle that allows it to function during neuroretina development, yet its absence from the ciliary basal body limits recruitment of a few TZ proteins likely underlying late-onset RP in both patients.

## Discussion

Human ciliopathies are a diverse group of syndromic and non-syndromic diseases that involve several organ systems as cilia are ubiquitous cellular organelles (18). Understanding how ciliary genes act in a tissue-specific manner to produce the wide variety of human phenotypes remains an active area of study. Here, we identified a homozygous frameshift variant in RP patients from 2 unrelated families. We find that *CEP162* is expressed in human retina and localizes to the basal body of outer segments in mice photoreceptors, and we provide strong functional evidence that *CEP162* plays a dual role in the retina.

In patient fibroblasts, low levels of *CEP162* mRNA were preserved and sufficient to produce an – albeit heavily truncated – CEP162 protein. We show that the residual truncated CEP162 protein maintains microtubule binding and is present at the mitotic spindles in patient-derived fibroblasts. These cells also had a normal complement of chromosomes and growth rates, consistent with previous *CEP162* knockdown in RPE1 cells (12). CEP162’s role in cell division was previously attributed to post-mitotic arrest of retinal neurons (7, 8). We found that knockdown of *CEP162* in developing mouse retina increased neuronal cell death, which was rescued by expression not only of full-length, but also truncated CEP162. This suggests that the truncated CEP162-E646R*5 protein retains function in neuroretina development likely due to its preserved microtubule binding activity. Mutations that disrupt CEP162 microtubule binding would likely result in defective retinal neurogenesis.

In quiescent cells, CEP162 is known to play a functional role in ciliogenesis by promoting TZ assembly (12). Truncated CEP162-E646R*5 protein was unable to localize to the ciliary basal body. The loss of CEP162 from the cilia results in reduced recruitment of some TZ proteins to cilia in patient fibroblasts. We confirm through rescue experiments that the truncated CEP162 is unable to restore the loss-of-function phenotype at the cilium. In the patient fibroblasts, we found that cilia formation was delayed due to the persistence of CP110 at the mother centriole. CP110 caps the distal end of centrioles, and its removal and degradation are prerequisites for ciliation. Other aspects of early ciliogenesis, like the acquisition of distal appendages and fusion of the ciliary vesicle to the plasma membrane, were unaffected in patient’s fibroblasts. Interestingly, after extended serum starvation, there was a dramatic increase in generation of – although abnormally long – cilia, showing that ciliogenesis does occur in the absence of CEP162.

It is interesting to speculate that perhaps the retinal defect in the patient (with residual CEP162 neurogenesis function) is not due to impaired ciliogenesis *per se*, as we find that cilia can still be formed, but instead due to reduced recruitment of TZ proteins, such as NPHP1, to the photoreceptor cilium. This could result in structural defects that are particularly detrimental to the outer segment, impairing photoreceptor *maintenance* rather than *formation* – clinically manifesting as late-onset retinitis pigmentosa and not as congenital retinal disease (as in *CEP290*-associated Leber congenital amaurosis (19)).

In summary, our data uncover a dual role for the centriolar protein, CEP162, in retinal neurons to ensure proper neurogenesis and ciliary TZ assembly. Our genomic, cell-based, and *in vivo* data show that although the mitotic function of CEP162 during neuronal development is maintained, specific loss of CEP162 function at the cilium results in a novel retinal ciliopathy in humans. These findings highlight the tissue-specific roles of ciliary proteins and may be instrumental for future studies exploring diverse ciliopathies.

## Methods

### Clinical assessment

Patients underwent ophthalmologic evaluation, including slit lamp examination, widefield color fundus photography and blue light autofluorescence (CLARUS 500; Carl Zeiss Meditec Inc., Dublin, CA), and spectral domain optical coherence tomography (Spectralis SDOCT, Heidelberg Engineering, Germany; PLEX Elite 9000; Carl Zeiss Meditec Inc.). Abdominal ultrasound (and additional CT in Patient 2) was conducted to exclude cysts of the kidneys and liver fibrosis.

### Genetic and genomic studies

#### Genomic testing

For the index patients of both families, Patients 1 and 2, genomic DNA (gDNA) was extracted from leukocytes, and pooled libraries were paired-end sequenced on an Illumina NextSeq500 (Illumina, San Diego, CA). Nucleotides were numbered with nucleotide ‘A’ of the ATG as ‘c.1’ (HGVS guidelines, http://www.hgvs.org). Maximum minor allele frequency (MAF) cut-off was 1%. Variants were checked for presence in ClinVar and HGMD databases (20, 21) and classified based on ACMG and ACGS guidelines (22–25). *Patient 1*. Targeted NGS (IDT xGen® Inherited Diseases Panel v1.0; IDT Integrated Technologies, Coralville, IA), WES (IDT xGen® Exome Research Panel v1.0) and variant assessment were conducted as reported previously (26). After targeted NGS, variant analysis was performed for 204 genes knowingly associated with IRD, including *RPGR*_*ORF15*_ (SeqNext module of SeqPilot software, JSI Medical Systems). Genes associated with the Human Phenotype Ontology (HPO) term “Retinal dystrophy” (HP:0000556) were searched for pathogenic homozygous or potentially compound-heterozygous variants (varSEAK Pilot 2.0.2, JSI Medical Systems). After WES, the CCG pipeline (27) and interface (Varbank 2.0) were used for data analysis as described (28, 29). ExAC and gnomAD databases (30) (as of 02/2021) were searched for candidate variants identified in homozygous state. *Patient 2*. WES was performed using the SureSelect XT Human All Exon V6 kits (Agilent, Santa Clara, CA, USA). CLC Genomics Workbench version 7.0.5 (CLCBio, Aarhus, Denmark) was used for read mapping against the human genome reference sequence (NCBI, GRCh37/hg19), removal of duplicate reads, coverage analysis, and quality-based variant calling, followed by further variant annotation using Alamut Batch (Interactive Biosoftware, Rouen, France). Variants were scored heterozygous or homozygous and assessed with our in-house variant filtering and visualization tool. 275 RetNet genes were assessed (version 4 of the RetNet panel), followed by assessment of homozygous variants in the exome. Copy number variations (CNVs) were assessed using ExomeDepth (v1.1.10) (31). *Segregation analysis and screening for the CEP162 variant c*.*1935dupA p*.*(E646R*5) in IRD patients*. Available members of Families 1 and 2 and 70 RP patients from North Africa, including 43 from Morocco, were tested for the *CEP162* variant c.1935dupA p.(E646R*5) by Sanger sequencing (primer sequences: Supplemental Table 1). Pseudonymized WES data from 1,184 IRD cases were also analyzed for presence of the variant (Ghent University Hospital).

#### Run of homozygosity (ROH) detection and haplotype analysis

*Patient 1*. ROH were identified using HomozygosityMapper applying default settings (32). ROH <3 kb adjacent to each other were merged manually, reducing ROH from 71 to 67. *Patient 2*. ROH were initially mapped by genome-wide single-nucleotide polymorphism (SNP) chip analysis (HumanCytoSNP-12 BeadChip platform, Illumina). ROH (>1 Mb) were identified using PLINK software (33) integrated in ViVar (34) and ranked according to length and number of consecutive homozygous SNPs. *Patient 1 and 2*. ROHs were determined with AutoMap (35) using VCF files from both patients (hg38). After identification of individual ROH, shared ROH (coordinate-wise) were determined (Figure 2C, Supplemental Table 1). To define the shared ROH on chromosome 6 and the common haplotype containing the *CEP162* variant, all WES variants on chromosome 6 were considered, irrespective of their zygosity (Figure 2C).

#### Fibroblast culture from skin biopsy

Fibroblasts were isolated from a dermal punch biopsy of the upper arm of Patient 1 as described (36). The skin biopsy was cut in 18-24 equally sized pieces and plated on a 6-well plate coated with 0,1% Gelatin. Fibroblasts migrated out of the skin biopsies after 7-10 days and were split on 2 75 cm^2^ flasks after 3-4 weeks. At 90% confluency, the 2 flasks were split on 3 175 cm_2_ flasks. After isolation, fibroblasts were routinely cultured in IMDM/Glutamax containing 15% fetal bovine serum (FBS) and 1% penicillin/streptomycin (all Invitrogen, Waltham, MA). Normal Primary Human Dermal Fibroblasts (HDFa, ATCC #PCS-201-012; Gaithersburg, MD) served as controls.

#### Karyotyping

30 metaphases from cultured Patient 1 fibroblasts were analyzed (standard procedures).

### Mouse studies

Mice protocols were approved by IACUC at the University of Michigan (registry number A3114-01). Albino Cetn2-GFP transgenic mice were obtained from Jackson Labs (Strain # 027967; Bar Harbor, ME). The *Pde6b*_*Rd1*_ mutation was removed by backcrossing to a BALB/cJ albino mouse from Jackson Labs. Albino CD-1 wild-type mice were from Charles River (Strain # 022; Mattawan, MI). All mice were housed under a 12/12-hour light cycle. Experimenters were not blinded to genotype. *In vivo* electroporation: Retinal transfection of CD-1 neonatal mice was conducted as previously described (17, 37, 38). In P0-P2 neonates, 2 μg/μl shRNA and 1 μg/μl rhodopsin-mCherry plasmid DNA was deposited subretinal. Retinal tissue was collected at P20-P22 for protein localization studies and collected at P14-15 for nuclear counts after shRNA knockdown.

### *In vitro* functional studies

#### DNA constructs

DNA constructs were generated using standard PCR-based subcloning methods. *Homo sapiens* centrosomal protein 162 (CEP162), transcript variant 1, mRNA (Accession: NM_014895.3) was obtained from GeneCoepia (HOC20483; Rockville, MD). FLAG-tag added by overlap extension PCR to N-terminus. All DNA constructs were cloned between a 5’ *Age*I and a 3’ *Not*I site, using standard T4 DNA ligation methods, and sequence confirmed. For mouse *in vivo* electroporation, the pRho plasmid was used (Addgene, plasmid # 11156; Watertown, MA). For mammalian cell culture, the pEGFP-N1 vector was used (Clontech, PT3027-5; Mountain View, CA). Mutagenesis was done using the QuikChange II XL kit (Stratagene, La Jolla, CA). Rho-mCherry was previously used (39). For primers, see Supplemental Table 2.

#### Plasmid expression in cell culture

AD293T or IMCD3 cells were transfected at 90-95% confluency with DNA constructs mentioned above, using Lipofectamine 3000 transfection reagent (Invitrogen, L3000001). The next day, cells were incubated in complete media or serum-free media for another 24 hours before analysis.

#### Immunofluorescence

Supplemental Table 3 contains a complete list of antibodies and dilutions. *Cetn2-GFP mouse retinal cross-sections:* Immunostaining was carried out as described (40). Briefly, fresh eyecups were embedded in OCT compound and quick-frozen in methylbutane cooled with liquid nitrogen. 10 μm thick retinal cryo-sections were blocked in 5% donkey serum and 0.05% Triton X-100 in PBS for 15 minutes before primary antibodies diluted in block for 1 hour. Sections were rinsed, fixed for 5 minutes in 1% paraformaldehyde in PBS, rinsed and incubated with secondary antibodies in block for 2 hours at 22ºC.

#### Electroporated mouse retinal cross-sections

The immunostaining protocol was adapted from Robichaux *et al*. (41). Electroporated retinas with mCherry expression were dissected in Supplemented Mouse Ringers, pH 7.4 and ~313-320 mOsM. Fresh retinas were blocked in 10% normal donkey serum, 0.3% saponin, 1× cOmplete_TM_ Protease (Millipore Sigma; Burlington, MA) diluted in Supplemented Mouse Ringers for 2 hours at 4°C. Primary antibodies diluted in block were incubated for 20-22 hours, rinsed and secondary antibodies incubated for 2 hours, all at 4°C. Retinas were rinsed, fixed in 4% paraformaldehyde for 30 minutes at 22°C, embedded in 4% agarose (Thermo Fisher Scientific, BP160-500), and 100 μm vibratome retinal-sections collected.

#### Cultured cells

50-60×10^3^ fibroblasts were plated onto 13 mm poly-L-lysine glass coverslips (Corning, 354085; Glendale, AZ) and grown overnight at 37ºC with 5% CO_2_. Serum-free DMEM for 24, 48 or 72 hours to induce ciliation. Cells were placed on ice for 10 minutes before fixation in 1% paraformaldehyde for 5 minutes followed by 15 minutes in ice-cold methanol. Cells were then rinsed, permeabilized for 5 minutes with 0.1% SDS in PBS, rinsed, and blocked with 0.1% Triton-X, 5% normal donkey serum and 5% BSA in PBS. Primary antibodies diluted in block were incubated overnight at 4ºC, rinsed, and incubated with secondary antibodies for 1 hour at 22ºC.

For all conditions, 10 μg/ml Hoechst 33342 was added with donkey secondary antibodies conjugated to Alexa Fluor 488, 568, or 647 (Jackson ImmunoResearch; West Grove, PA). Coverslips (#1.5, Electron Microscopy Sciences; Hatfield, PA) were mounted with Prolong Glass (Thermo Fisher Scientific). Images acquired using a Zeiss Observer 7 inverted microscope with a 63× oil-immersion objective (1.40 NA), LSM 800 confocal with Airyscan detector controlled by Zen 5.0 (Carl Zeiss Microscopy; White Plains, NY). Image manipulation was limited to adjusting the brightness level, image size, rotation and cropping using FIJI (ImageJ, https://imagej.net/Fiji).

#### Immunoprecipitation

A 10 cm plate of AD293T cells was transfected using Polyethyleneimine (PEI, Sigma 408727) at 6:1 ratio PEI:DNA with 10 μg of mock, FLAG-CEP162, or FLAG-CEP162-E646R*5 construct. After 24 hours, cells were lysed in 1% NP40 in PBS, cleared by centrifugation at 15,000 rpm for 20 min at 10°C and supernatant incubated on anti-FLAG M2 magnetic beads (Millipore Sigma, M8823) rotating overnight at 4°C. Immunoprecipitants were eluted with 100 mM 3X FLAG peptide (Millipore Sigma, F4799) in lysis buffer rotating for 1 hour at 4°C and processed for immunoblotting.

#### Microtubule pelleting assays

FLAG-CEP162 or FLAG-CEP162-E646R*5 immunoprecipitated in BRB80 (80 mM PIPES, 1 mM MgCl_2_, 1 mM EGTA, ph 6.8, diluted from a 5X stock with 1x Protease Inhibitor) from 5 10 cm plates of AD293T cells was added to taxol-stabilize microtubules. The binding reaction was added on top of a 1 mL cushion (40% glycerol in BRB80 with 10 μM taxol, 1x Protease Inhibitor and 1 mM GTP) and spun in fixed-angle rotor TLA120.1 at 76,900 rpm for 40 minutes at 25°C. Soluble and pellet fractions were collected, run on SDS-PAGE, and Western blotted for FLAG.

### Statistics

All data represent at least 3 independent experiments. The data are presented as a mean +/-SD or SEM as indicated in the figure legends. A Welch’s t test was performed using Prism9 (GraphPad) with corresponding P values listed in the legends.

### Study approval

All individuals involved gave their informed consent prior to inclusion in this study. All investigations were conducted according to the Declaration of Helsinki, and the study was approved by the Ethics Committee of the University Hospital of Cologne, Ghent (EC UZG 2017/1540) and Brussels (A420701EI13L).

## Author contributions

NN: Project design, acquisition of data, analysis and interpretation of data, drafting and revising the manuscript.

KVS: Acquisition of data, analysis and interpretation of data, revising the manuscript. SL: Acquisition of data, analysis and interpretation of data.

MBW: Acquisition of data, analysis and interpretation of data, drafting the manuscript. ADR: Acquisition of data, analysis and interpretation of data, drafting the manuscript. SK: Acquisition of data, analysis and interpretation of data.

MJ: Acquisition of data, analysis and interpretation of data.

JRW: Acquisition of data, analysis and interpretation of data, revising the manuscript. JW: Acquisition of data, analysis and interpretation of data.

HT: Acquisition of data, revising the manuscript.

NG: Acquisition of data, analysis and interpretation of data.

MVH: Acquisition of data, analysis and interpretation of data.

JW: Acquisition of data, analysis and interpretation of data.

RM: Acquisition of data, analysis and interpretation of data.

SGD: Acquisition of data, analysis and interpretation of data.

JVD: Acquisition of data, analysis and interpretation of data.

AH: Acquisition of data, analysis and interpretation of data.

HS: Acquisition of data, analysis and interpretation of data.

LM: Acquisition of data, analysis and interpretation of data.

AFR: Acquisition of data, analysis and interpretation of data.

TL: Acquisition of data, analysis and interpretation of data.

KD: Acquisition of data, analysis and interpretation of data.

DR: Acquisition of data, analysis and interpretation of data.

KMW: Acquisition of data, analysis and interpretation of data.

MVL: Acquisition of data, analysis and interpretation of data.

HR: Acquisition of data, analysis and interpretation of data.

FH: Acquisition of data, analysis and interpretation of data.

PN: Acquisition of data, analysis and interpretation of data.

HTH: Acquisition of data, analysis and interpretation of data.

UZ: Conception and project design, acquisition of data, analysis and interpretation of data, drafting and revising the manuscript.

JNP: Conception and project design, acquisition of data, analysis and interpretation of data, drafting and revising the manuscript.

EDB: Conception and project design, acquisition of data, analysis and interpretation of data, drafting and revising the manuscript.

HJB: Conception and project design, acquisition of data, analysis and interpretation of data, drafting and revising the manuscript.

## Acknowledgements

We are grateful to Woo Jung Cho for assisting the design of CLEM experiments, Dr. Ryoma Ohi for reagents and guidance on the microtubule pelleting assays, and Dr. Kristen Verhey for expert feedback on the manuscript. This work was supported by a NIH P30 grant EY007003 (University of Michigan), NIH T32 grant HD007505 (NN), Matilda E. Ziegler Research Award (JNP), Career Development Award (JNP), an Unrestricted Grant (University of Michigan) from Research to Prevent Blindness, and grants from PRO RETINA Deutschland, from Stiftung Auge (Deutsche Ophthalmologische Gesellschaft; HJB), from Dr. Senckenbergische Stiftung and by Köln Fortune (University Hospital of Cologne; HJB), from Ghent University Special Research Fund (BOF20/GOA/023) (EDB); Ghent University Hospital Innovation Fund NucleUZ (EDB); John W. Mouton Pro Retina Fund (EDB); H2020 MSCA ITN grant (No. 813490 StarT) (EDB), EJPRD19-234 Solve-RET (EDB). EDB is a senior clinical investigator (1802220N) and KVS was a postdoctoral fellow (12W7218N) of the Research Foundation-Flanders (FWO); ADR is Early Starting Researcher of StarT (grant No. 813490). EDB is member of ERN-EYE (Framework Partnership Agreement No 739534-ERN-EYE).

## SUPPLEMENTARY MATERIAL

**SUPPLEMENTAL TABLE 1**

**AutoMap output for Patient 1 and Patient 2**.

**SUPPLEMENTAL TABLE 2**

**List of primers** used for PCR amplification and Sanger sequencing of exon 15 of *CEP162*, for qRT-PCR experiments and for the generation and sequencing verification of FLAG-tagged CEP162 plasmids.

**SUPPLEMENTAL TABLE 3**

**List of antibodies** used in this study including the source, catalog number, and dilution used in either immunofluorescence (IF) or Western blot (WB) experiments.

**SUPPLEMENTAL FIGURE 1**

**Abdominal CT of Patient 2. A** Kidneys (white arrows) without cysts or other morphological abnormalities. **B** Pancreas (grey arrow) with a lipomatous aspect.

**SUPPLEMENTAL FIGURE 2**

**Homozygous *CEP162* frameshift variant in Patient 2 and segregation analysis in his daughter. A** Whole-exome sequencing reads visualized in the Integrative Genomics Viewer IGV, showing the homozygous *CEP162* variant c.1935dupA (p.(E646R*5)). **B** Sanger sequencing electropherogram of the heterozygous *CEP162* variant in the daughter of Patient 2.

**SUPPLEMENTAL FIGURE 3**

***CEP162 mRNA* expression is reduced, but truncated CEP162-E646R*5 protein is expressed in patient fibroblasts. A** Transcript levels of *CEP162* in control and patient fibroblasts after 24 hours with (+) or without (–) serum conditions. Quantitative RT-PCR using primer sets across exons 5, 14 and 25 of the *CEP162* gene was performed on 4 replicates at each condition. The relative mRNA expression level in each sample was normalized to the housekeeping gene *HSP90*. Error bars represent SD. *p<0.0459, **p<0.0012. **B** Inhibition of NMD by anisomycin treatment results in a similar increase in fibroblasts of patient and controls. Bar and scatter plot of 3 independent experiments analyzed in duplicates by ddPCR. **C-D** Representative Western blot of control and patient fibroblast lysates. Blots were probed with an N-terminal anti-CEP162 antibody to detect full length (red arrow) and truncated (red asterisk) bands (**C**). C-terminal anti-CEP162 antibody only detected full length CEP162 (red arrow) in the control fibroblast lysates. α-tubulin was used as a loading control.

**SUPPLEMENTAL FIGURE 4**

**FLAG-CEP162 and FLAG-CEP-E646R*5 expression in 293T cells and glutamylated tubulin staining in control and patient fibroblasts. A** Representative Western blot of 239T cell lysates from untransfected control, FLAG-CEP162 (~165 kDa, red arrow) and FLAG-CEP162-E646R*5 (~80 kDa, red asterisk). Blots were probed for FLAG to detect expressed protein as well as α-Tubulin as a loading control. **B** Control and patient fibroblasts with and without a cilium were co-stained with Arl13b and glutamylated tubulin B3 (GlutTub, magenta). Nuclei labeled with Hoechst 33342 (grey). Scale bars, 5 μm and 2 μm. Bar graph quantifying the % of control or patient fibroblasts with positive glutamylated tubulin staining shown on the right. *p<0.011

**SUPPLEMENTAL FIGURE 5**

**Quantification of CEP290, CP110 and RP2 protein levels in CEP162-mutant patient fibroblasts compared to controls**. Representative Western blots show serial dilutions of control (CEP162-WT) fibroblast and patient (CEP162-E646R*5) fibroblast lysates for CEP290, CP110 and RP2 proteins. Amount of protein lysate (μg) loaded in each lane is displayed below. The fluorescent signal produced by the CEP290 or CP110 bands in 3 separate experiments was plotted versus total protein loaded. The slope of the curves was used to calculate the amount of each protein in control and patient fibroblasts.

**SUPPLEMENTAL FIGURE 6**

**Immunostaining for proteins implicated in early stages of ciliogenesis and removal of CP110 in control and patient fibroblasts. A** Immunostaining of early ciliogenesis markers in control and patient fibroblasts: EDH1, CEP164, and IFT88 (magenta). **B** Molecules implicated in the removal of CP110: TTBK2 and MPP9 (magenta) are normally localized at the basal body of patient fibroblasts.

**SUPPLEMENTAL FIGURE 7**

**Expression analysis of CEP162. A** Mining of single-cell transcriptional retinal data displaying the highest expression in cells belonging to the ganglion cell population. (Left) UMAP plot of 19,768 human neural retinal cells colored based on cell type. Visualization of scaled and corrected expression on UMAP plot for CEP162. (Top right) Violin plot for visualization of scaled and corrected expression of *CEP16*2 in the 9,768 human neural retinal cells. (Bottom right) Individual dots are colored and sized to visualize both the proportion of cells in each population expressing a given gene (dot diameter size) and the average expression level of that gene (dot color intensity) in the detected cells. **B-F** CEP162 immunostaining on human retinal sections. Representative fluorescence images of human paraffin-embedded sections stained with anti-CEP162 antibody (red) (**B-C**) or negative control (**D-E**). Sections were counter-stained with DAPI (blue), displayed in **C** and **E**. Scale bars represent 100 mm. **F** Immunostaining of the CEP162 protein on human retina is visible in light brown, displaying retinal expression. Light microscopy at 400x. Scale bars represent 100 mm. Abbreviations: GCL, ganglion cell layer; INL, inner nuclear layer; IPL, inner plexiform layer; OPL, outer plexiform layer; ONL, outer nuclear layer; OLM, outer limiting membrane; PR, photoreceptors.

**SUPPLEMENTAL FIGURE 8**

**Uncropped Western blot images**. Red boxes delineate the cropped area of the blot shown in the indicated figures.

